# Novel functions of Kaposi’s sarcoma herpesvirus viral FLICE inhibitory protein

**DOI:** 10.1101/629113

**Authors:** Kevin Herold, Ayana Ruffin, Mathew Constantine, Amerria Causey, Ashley E. Mitchell, Jennifer C. Chmura, Razia Moorad, Elana S. Ehrlich

**Affiliations:** Department of Biological Sciences, Towson University, Towson, MD, USA; Lineberger Comprehensive Cancer Center, School of Medicine, Department of Immunology and Microbiology, The University of North Carolina at Chapel Hill, Chapel Hill, NC, USA

## Abstract

KSHV viral FLICE inhibitory protein (vFLIP) is a potent activator of NF-κB signaling and an inhibitor of apoptosis and autophagy. Inhibition of vFLIP function and NF-κB signaling promotes lytic reactivation. Here we provide evidence for a novel function of vFLIP in promoting NF*κ*B signaling through inhibition of the DUB activity of the negative regulator, A20. We demonstrate interaction of vFLIP with the Itch/A20 ubiquitin editing complex. We have identified a SUMO interaction motif in vFLIP that is required for NF-κB activation. Mutation of the SIM in BAC16 resulted in increased spontaneous RTA expression and loss of spindle cell morphology. Our results suggest a role for SUMO in mediating vFLIP function and provide evidence for vFLIP modulation of the negative regulation of NF-κB signaling by A20. Our results provide further insight into the function of vFLIP and SUMO in the regulation of NF-κB signaling and the latent lytic transition.

## Introduction

Kaposi’s sarcoma herpesvirus is a member of the γ2 subfamily of herpesviruses and the causative agent of Kaposi’s sarcoma[1,2]. The KSHV genome has been found in the cells of two B-cell lymphoproliferative diseases: primary effusion lymphoma (PEL) and Multicentric Castleman’s disease (MCD) and is associated with two inflammatory syndromes, immune reconstitution inflammatory syndrome-KS (IRIS-KS) and KSHV inflammatory cytokine syndrome (KICS)[3–5]. KSHV has been classified as a Group 1 carcinogen by the International Agency for Research on Cancer and the National Toxicology Program 14^th^ Report on Carcinogens[6].

The KSHV genome contains several viral homologs of cellular genes, many of which promote immune evasion, cell survival and proliferation. KSHV exists mostly as a latent infection, where the viral genome is tethered to the host chromosome by LANA and infectious virions are not produced. Nascent virions are produced during periods of lytic replication induced by expression of the viral transactivator, RTA[7].

KSHV oncogenesis is, in part, attributed to genes expressed during latency. Viral FLICE inhibitory protein (vFLIP or K13), is a latently expressed gene that was originally identified as an inhibitor of apoptosis, due to the presence of tandem death effector domains[8,9]. vFLIP is a potent activator of NF-κB signaling and this activity is dependent on interaction with IKK*γ* [10–12]. vFLIP has also been shown to promote NF-κB signaling through upregulation of IKK*ε* and CADM1 and inhibition of the SAP18/HDAC1 complex resulting in activation of NF-κB via acetylation of p65[13–15]. NF- κB signaling is required for the virus to maintain latency, as chemical inhibition of this signaling pathway has been shown to promote lytic replication[16,17].

vFLIP plays a role in oncogenesis and genome instability. A transgenic mouse model of vFLIP expression displays persistent NF-κB activation and an increased incidence of lymphoma as well as B cell abnormalities similar to those observed in MCD (reviewed in [18]). More recently, vFLIP was shown to increase LINE-1 retrotransposition which may promote genome instability[19].

NF-κB signaling induces expression of negative regulators that limit the inflammatory response. A20 (TNFAIP3), one such negative regulator of NF-κB, is induced by vFLIP. A20 is a ubiquitin editing protein with both C-terminal ubiquitin ligase activity and N-terminal deubiquitinase activity. In one well characterized mechanism, A20 forms a ubiquitin editing complex with Itch, RNF11, and TAX1BP1, and downmodulates NF-κB signaling through removal of K63-linked polyubiquitin chains from RIPK1 followed by addition of K48-linked polyubiquitin chains, resulting in degradation of RIPK1 via the proteasome. A20 is reported to deubiquitinate a number of signaling intermediates within the NF-κB pathway in addition to RIPK1, including IKK*γ*, TRAF6, TRAF2 and MALT1 [20–22].

Here we report two observations suggesting novel functions of vFLIP. 1. vFLIP inhibition of A20 activity and 2. vFLIP has a SUMO interaction motif (SIM) that functions in maintaining latency. We previously reported that RTA induces the degradation of vFLIP early in lytic reactivation resulting in the termination of NF-κB signaling, presumably to promote transition from latency to lytic replication[23]. RTA induced degradation of vFLIP is dependent on the activity of the Itch ubiquitin ligase[24]. We identified mutants of vFLIP that are unable to interact with Itch and cannot activate NF-κB[24]. Here we report that vFLIP interacts with the A20/Itch ubiquitin editing complex and this interaction occurs independently of RTA. We propose that vFLIP inhibits A20 DUB activity to modulate NF-κB signaling through interference with negative regulation. We demonstrate reduced A20 activity and increased levels of RIPK1K ubiquitin conjugates following stimulation with TNFA in the presence of vFLIP.

Small Ubiquitin-like Modifier (SUMO) proteins, when covalently conjugated to substrate proteins, can modulate the stability, interaction and activity of proteins. We have identified a SUMO interacting motif (SIM) in vFLIP. vFLIP SIM mutants exhibited reduced SUMO1 and SUMO 2/3 binding and were unable to activate NF-κB signaling. Small molecule inhibition of SUMO conjugation resulted in increased virus production, suggesting a role for SUMO the latent to lytic transition. We generated a BAC16 mutant where the V22 in the vFLIP SIM is changed to E. iSLK cells containing the vFLIP SIM mutant exhibited higher levels of spontaneous reactivation and lost their spindle shaped morphology. Taken together, our findings suggest a novel role for vFLIP in activation of NF-κB signaling via inhibition of the DUB activity of A20 and that a SIM in vFLIP plays a role in maintaining latency. Future studies will need to determine whether the SIM in vFLIP is required for interaction with A20 and whether this interaction is required for maintaining latency. These observations expand our understanding of vFLIP function and have the potential to increase our understanding of the mechanisms governing the maintenance of latency.

## Results and Discussion

### vFLIP interacts with Itch and A20

We previously reported that RTA induces the degradation of vFLIP via the cellular ubiquitin ligase, Itch. We hypothesized that RTA was recruiting Itch to vFLIP to promote ubiquitination and degradation of the viral protein. Upon further characterization of the interactions between vFLIP and Itch as part of the Itch/A20 ubiquitin editing complex, we observed interaction between vFLIP, Itch and A20, even in the absence of RTA (Fig 1a-b). Based on this data we hypothesized that vFLIP interacts with the Itch/A20 ubiquitin editing complex in latency and RTA expression, occurring early in lytic reactivation, results in activation of Itch and/or A20 ubiquitin ligase activity (either directly or indirectly), resulting in the subsequent ubiquitination and degradation of vFLIP. If this model were correct, we would expect to see degradation of additional targets of Itch and/or A20 in the presence of RTA. In fact, we detected a modest decrease in the levels of A20, a known target of Itch, in cells transfected with RTA compared to empty vector control (Fig. 2a). We evaluated additional Itch and/or A20 substrates, RIPK1 and c-Jun, by immunoblot and observed a modest decrease in protein levels compared to the internal tubulin control (Fig. 2b).

**Figure 1.**
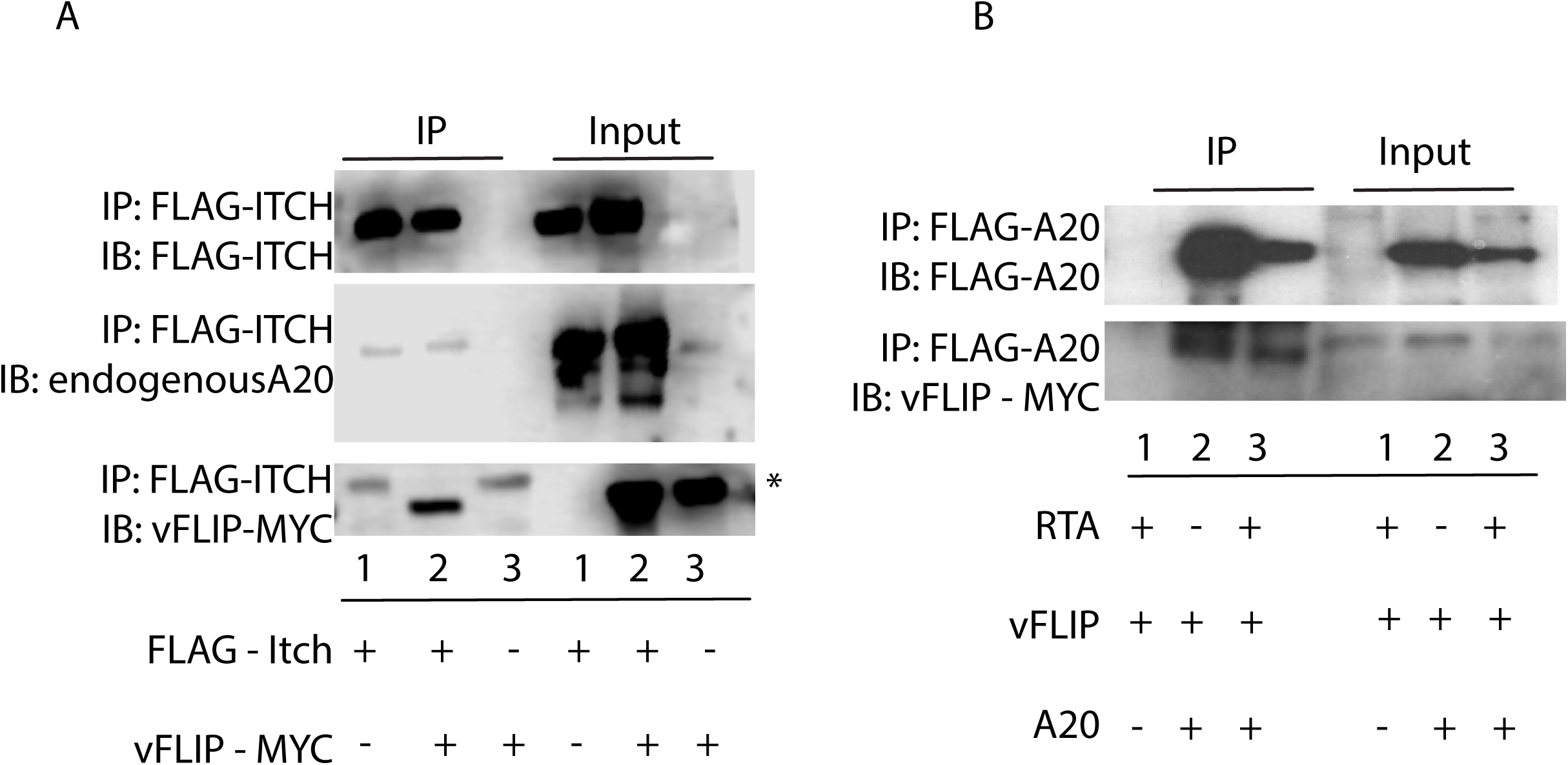
vFLIP interacts with Itch and A20. A. vFLIP does not interfere with Itch/A20 interaction. 293T cells were transfected with Flag tagged Itch and/or myc tagged vFLIP where indicated. Itch was immunoprecipitated with Flag antibody and immunoprecipitates were analyzed by immunoblot against Flag, myc, and endogenous A20. B. vFLIP interaction with A20 occurs independent of RTA. 293T cells were transfected with Flag tagged A20, myc tagged vFLIP and/or RTA where indicated. Itch was immunoprecipitated with Flag antibody and immunoprecipitates were analyzed by immunoblot against Flag and myc.

**Figure 2.**
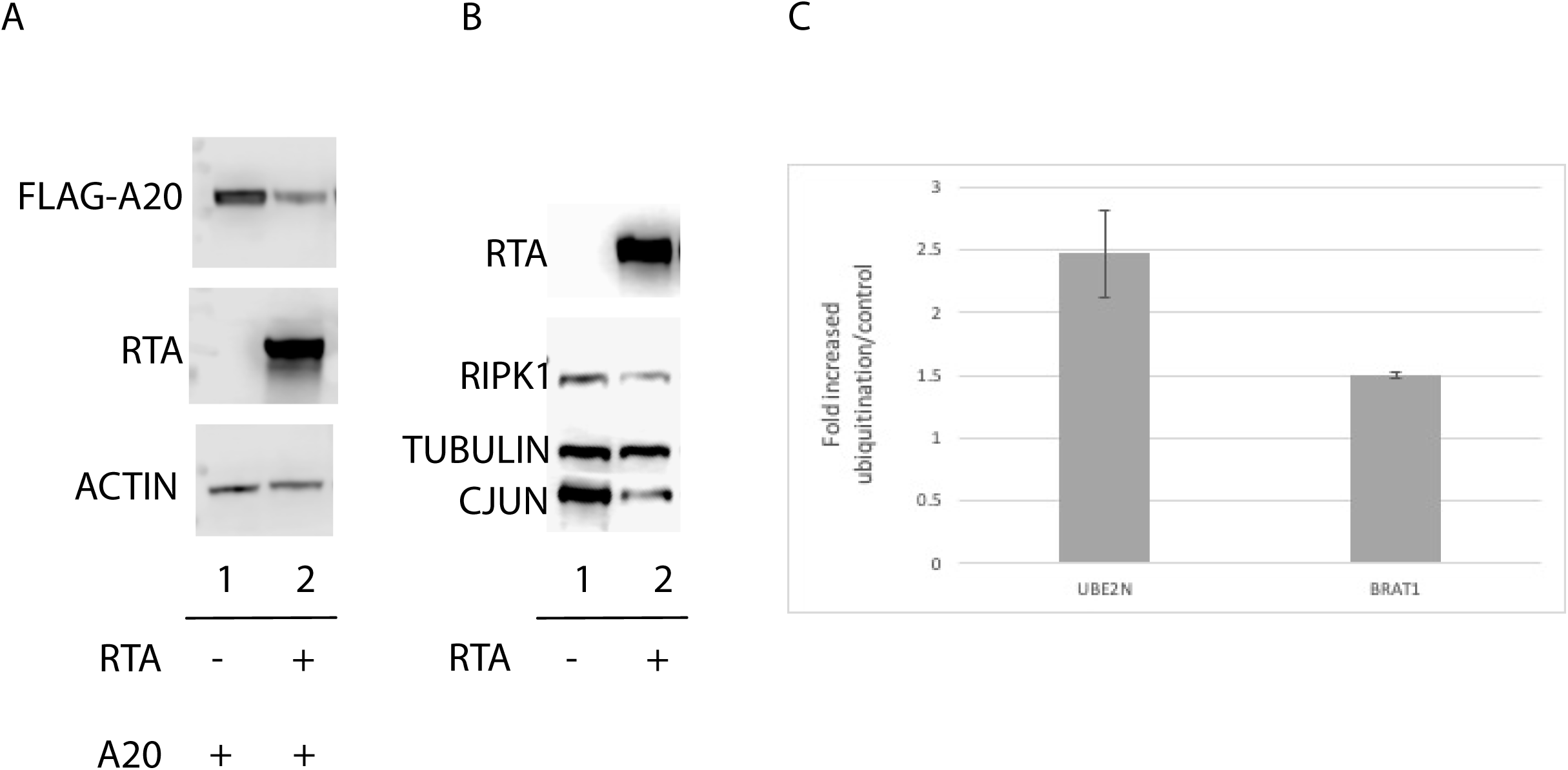
RTA expression is associated with reduced stability of Itch substrates. A. A20, and B. cJun, and RIPK1 protein levels are reduced in the presence of RTA. 293T cells were transfected with Flag tagged A20, RTA or empty vector where indicated. Cell lysates were probed for Flag or endogenous cJun or RIPK. Actin or tubulin was used as a loading control. C. RTA expression induces alterations in the ubiquitinated proteome. 193 sites in 146 proteins displayed differential ubiquitination in RTA expressing cells compared to vector transfected controls. SILAC labeled 293T cells were transfected with RTA or empty vector control. Lysates were processed for mass spectrometry and ubiquitinated peptides were enriched using the anti-diglycine remnant (K-ε-GG) antibody. Samples were cleaned on C18 and analyzed by MS. MS spectra were processed and analyzed by Maxquant and Perseus. D. UBE2N and BRAT 1, known substrates of A20 and Itch displayed increased ubiquitination in RTA transfected cells.

To further examine the effect of RTA on the ubiquitination of Itch and A20 substrates, we conducted an analysis of the ubiquitinated proteome in SILAC labeled RTA transfected 293T cells using the anti-diglycine remnant (K-ε-GG) antibody. We detected two known Itch or A20 substrates that exhibited a significant increase in ubiquitination in the presence of RTA. We observed a 1.5-fold increase in the ubiquitination of BRAT1, a known Itch substrate, and a nearly 3-fold increase in ubiquitinated UBE2N, a known A20 substrate, in RTA transfected cells compared to empty vector transfected controls (Fig. 2c). Taken together, this data suggests that while RTA may play a role in the activation of the Itch ubiquitin ligase or Itch/A20 ubiquitin editing complex, the exact mechanism remains unclear.

### vFLIP inhibits the debiquitinase activity of A20

A20 is a well characterized negative regulator of NF-κB signaling. Following stimulation of NF-κB via the TNF receptor (TNFR), A20 downregulates signaling by removal of K63 linked polyubiquitin chains from RIPK1 and in concert with Itch, adds K48 linked polyubiquitin, resulting in RIPK1 degradation via the proteasome. It was previously reported that vFLIP induces the expression of A20. It has been proposed that A20 expression, in the context of latent KSHV infection, is necessary to limit the inflammatory phenotype induced by persistent NF-κB signaling. We hypothesized that A20 activity needs to be tightly regulated, as excessive activity has the potential interfere with latency and cell survival, and inhibition of NF-κB signaling has been shown to promote apoptosis and lytic reactivation. To this end, we assessed the impact of vFLIP on A20 DUB activity. Using purified K63-linked tetraubiquitin, A20 and vFLIP, we evaluated A20 DUB activity via *in vitro* assay. Addition of purified A20 alone to tetraubiquitin, resulted in cleavage of tetraubiquitin to faster migrating mono and polyubiquitin species (Ub-3, Ub-2, Ub-1), however addition of recombinant vFLIP resulted in a dose dependent decrease in DUB activity (Fig. 3a). A well characterized target of A20 DUB activity is RIPK1, following TNFR stimulation. Within 30 minutes of TNFR stimulation, transient K63-polyubiquitin conjugates of RIPK1 can be detected via western blot. By 2hrs post stimulation, K63 ubiquitin conjugates are removed by A20. To determine whether vFLIP inhibits DUB activity in the context of NF-κB signaling, we evaluated K63 linked RIPK1 ubiquitin conjugates following stimulation with TNFA. Cells were transfected with wild-type or K63-only ubiquitin and vFLIP where indicated. Endogenous RIPK1 was purified and immunoprecipitates were probed for HA-tagged wild type or K63 only ubiquitin. Control cells, lacking vFLIP, displayed the characteristic increase in RIPK1 ubiquitin conjugates after 30 minutes of TNFA treatment, followed by deubiquitination 2h post treatment (Fig. 3b). Addition of vFLIP, however, resulted in detection of sustained RIPK1 ubiquitin conjugates, which was accentuated in cells transfected with K63-only ubiquitin (Fig. 3b). These data, taken together, suggest that vFLIP has an inhibitory effect on A20 DUB activity.

**Figure 3.**
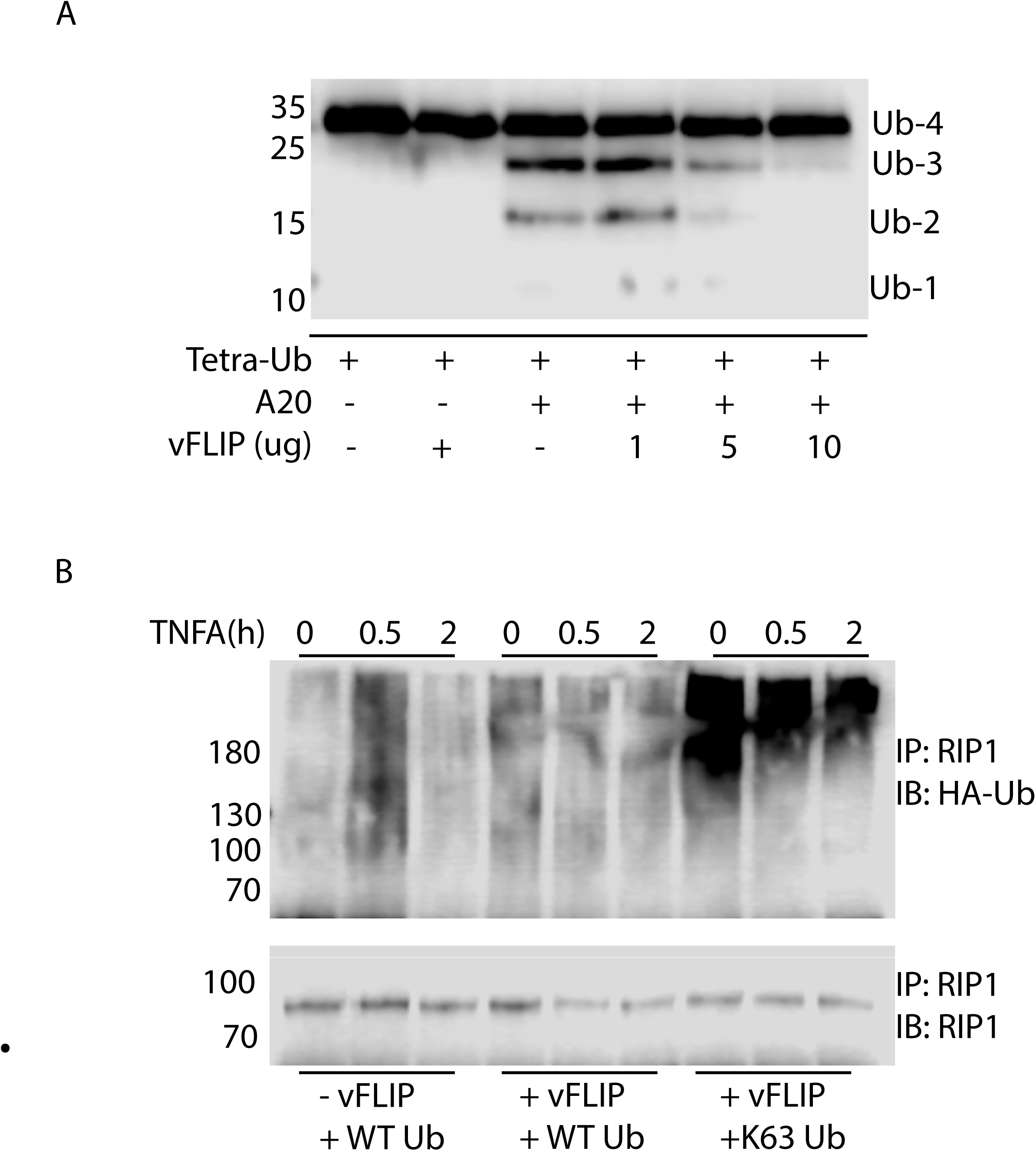
vFLIP inhibits A20 deubiquitinase activity. A.vFLIP inhibits A20 deubiquitinase activity *in vitro*. In 20μl reactions the following reagents were added where indicated: tetra-K63 Ub (Boston Biochem) (500nM), V5-vFLIP (1,5,10μM), and A20 (2μM). Reactions were incubated at 37ºC for 2hours following addition of 4x Laemmli loading buffer. Samples were run on 15% SDS- PAGE gel and analyzed via immunoblot with antibody against ubiquitin. B. vFLIP abrogates the deubiquitination of RIPK1. HEK 293T cells were transfected with either empty vector control, HA- WT Ub, and or myc-vFLIP, then treated with TNF at 0, 0.5, and 2 hours after transfection and before harvesting, and analyzed via immunoprecipitation with anti-RIP1. Following immunoprecipitation, lysates were analyzed by immunoblotting with anti-HA and anti-RIP1.

### vFLIP has a SUMO interaction motif (SIM)

PML nuclear bodies have been implicated in antiviral defense, apoptosis, and the DNA damage response as well as other cellular processes. They are known to be highly modified by the small ubiquitin-like modifier SUMO. In our initial vFLIP studies, we observed vFLIP colocalization with PML via immunofluorescence assay, prompting the question as to whether vFLIP has a SIM (unpublished observation). Analysis of the vFLIP protein sequence resulted in the identification of a putative tandem SIM spanning amino acids V21-VLFLLN-V28 based on the following consensus motif (V/L/I, X, V/L/I, V/L/I) or (V/L/I, V/L/I, X, V/L/I) (Fig. 4a). Indeed, wild-type vFLIP was able to bind recombinant SUMO 1 and SUMO 2/3, albeit to a lesser extent, while mutant vFLIP V22E, was no longer able to bind to either SUMO 1 or 2/3 (Fig. 4b).

**Figure 4.**
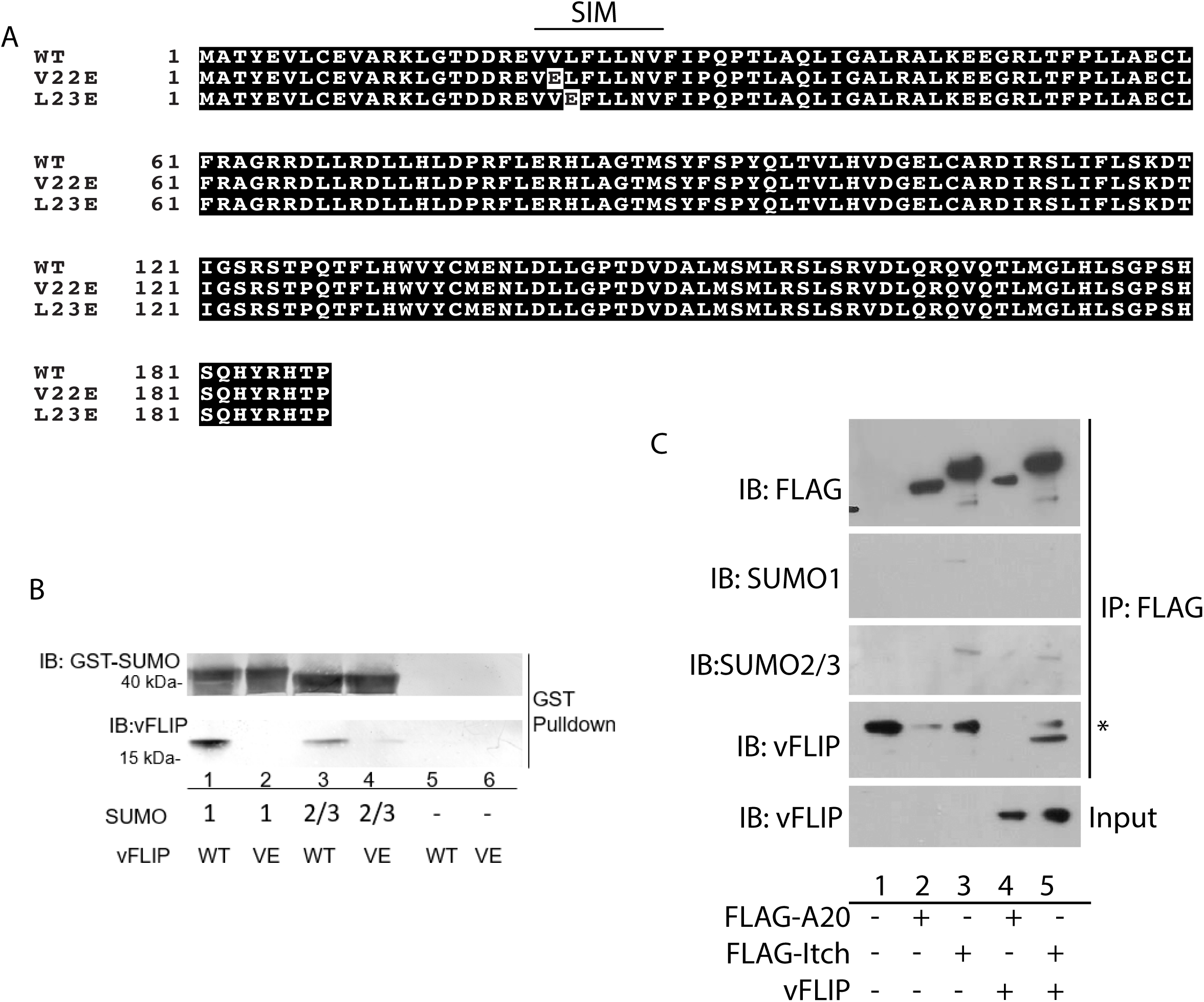
vFLIP has a SIM. A. Sequence alignment illustrating SIM location and mutations generated B. SIM dependent interaction between vFLIP and SUMO 1 and SUMO 2/3. Recombinant GST-SUMO was expressed in E. coli and 93T cells were transfected with myc- vFLIP. Interaction was evaluated via GST pull-down assay. Cell lysates containing myc tagged vFLIP were incubated with GST tagged SUMO bound to glutathione-sepharose. Purified proteins were analyzed via western blot. C. Evidence for Itch SUMOylation. 293T cells were transfected with Flag-Itch, Flag-A20 or empty vector where indicated. Cell lysates were processed for immunoprecipitation using anti-FLAG antibody. Immunoprecipitates were washed with a 500mM NaCl followed by analysis via SDS-PAGE and western blot against Flag or endogenous SUMO 1 or SUMO 2/3.

We previously reported that this vFLIP mutant was resistant to RTA induced degradation and was unable to activate NF-κB, suggesting that this SIM is important for interaction with components of the NF-κB signaling pathway and RTA[23,24]. vFLIP V22E is also unable to interact with the Itch ubiquitin ligase[24]. To determine whether Itch and A20 are SUMOylated, we immunoprecipitated FLAG tagged Itch or A20 and probed the 500mM NaCl washed immunoprecipitates with antibody against endogenous SUMO 1 or SUMO 2/3. We observed corresponding SUMO 1 and 2/3 bands in the Itch immunoprecipitates (Fig. 4c). Itch is not reported to be SUMOylated in the literature or in the Phosphosite plus database suggesting that either the modification is highly transient or that Itch is interacting with a SUMOylated protein. Taken together, these results support a model where vFLIP may interact with the Itch/A20 ubiquitin editing complex either directly or indirectly via a SUMO dependent mechanism.

### Mutation of the SIM in BAC16 results in loss of spindle cell morphology and increased RTA expression

Our observations suggest that SUMOylation plays a role in regulation of NF-κB signaling induced by vFLIP. Previous reports demonstrated the importance of vFLIP induced NF-κB signaling for maintaining viral latency and cell survival, as inhibition with Bay 11-7082 and transfection with an IKBA dominant negative mutant resulted in increased apoptosis and lytic reactivation[25][26]. We hypothesized that if SUMOylation was required for maintaining latency, we would see a dose dependent increase in lytic reactivation upon SUMOylation inhibition and in fact we did observe this in Vero rKSHV.294 cells treated with the small molecule SUMOylation inhibitor 2-D08 [27](manuscript in preparation).

To further explore the function of the vFLIP SIM, we constructed a V22E vFLIP mutant in BAC16 and generated iSLK BAC16 vFLIPV22E cells. The BAC16 vFLIPV22E mutant was sequenced and analyzed by digest with Sbf1 followed by pulsed field gel electrophoresis (PFGE). The BAC16 vFLIP V22E sequence was consistent with the BAC16 consensus except for the mutation we introduced into the vFLIP coding sequence (Fig. 5a). Through PFGE analysis, we observed similar banding patterns between wild type and mutant BAC16 except for a shift in the bands above 48kb suggesting a possible alteration in the terminal repeat length (Fig. 5b). We hypothesized that if the SIM in vFLIP is required for vFLIP function we should see an increase in spontaneous lytic reactivation. To evaluate lytic reactivation, we analyzed RTA expression in iSLK cells containing wild-type or mutant BAC16. As expected, doxycycline treatment resulted in a 9- fold increase in RTA expression in iSLK cells containing wild-type BAC16, however cells containing BAC16 vFLIPV22E displayed a 12-fold increase in RTA expression in the absence of doxycycline treatment, suggesting an increase in spontaneous lytic reactivation (Fig. 5c). Doxycycline induction of RTA expression was 2-fold higher in cells containing the SIM mutant compared to wild-type BAC16 (Fig5c, compare WT +DOX to VE +DOX). We observed a corresponding increase in RTA protein levels (Fig. 5d). Interestingly, we observed a striking difference in cell morphology when comparing iSLK cells containing wild-type and SIM mutant BAC16. Cells containing wild-type BAC16 had the characteristic spindle shape morphology associated with vFLIP induced activation of NF-ΚB, whereas in cells containing the SIM mutant virus, vFLIP had more of a cobblestone appearance, similar to iSLK cells lacking BAC16 (Fig 5e). Taken together these data support a functional role for the SIM in vFLIP, as virus lacking an intact SIM exhibited higher levels of spontaneous lytic reactivation and infected cells lost the characteristic vFLIP induced NF-κB associated spindle cell morphology.

**Figure 5.**
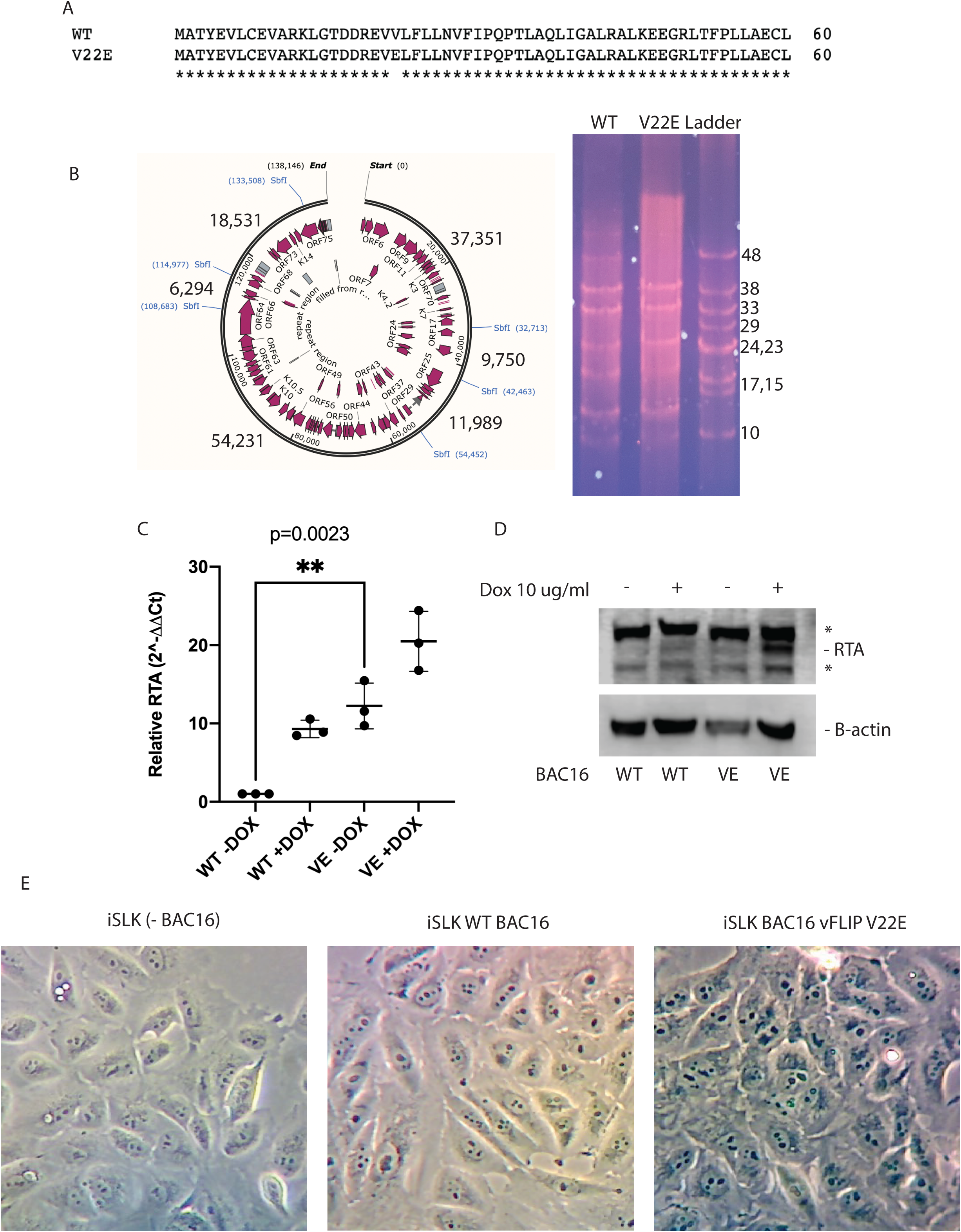
Disruption of the SIM in vFLIP results in increased lytic reactivation and loss of spindle shape morphology. A. Sequencing of BAC16 vFLIP V22E confirms that the BAC maintained the consensus sequence with the exception of the point mutation introduced into vFLIP. B. PFGE analysis of wild type and vFLIP V22E mutant BAC16. Wild-type and mutant BAC16 were digested with SbfI for 40min and separated on a 1% PFGE-grade agarose gel at 14 degrees C, for 8.5h at 6V/cm with initial and final switch times of 1 and 5 seconds, respectively. C. Spontaneous RTA expression is significantly increased in iSLK cells containing BAC16 vFLIP V22E compared to wild type BAC16. RTA expression was quantified in WT or SIM mutant BAC16 iSLK cells -/+ doxycycline treatment and normalized to RPS13 housekeeping gene. D. RTA expression is elevated in iSLK cells containing BAC16 vFLIP V22E compared to wild type BAC16. iSLK cells containing wild type or SIM mutant BAC16 were treated with doxycycline for 24h as indicated. Cells were washed with PBS and lysed directly in Laemmli Buffer. Lysates were analyzed by SDS PAGE followed by immunoblot against RTA and BACTIN as a loading control. * Indicate nonspecific bands. E. iSLK cells containing BAC16 vFLIP V22E lack vFLIP dependent spindle shape morphology. Live iSLK cells were imaged using a Celestron digital microscope imager and a Zeiss Primo Vert inverted microscope using a 10x objective.

## Discussion

We have presented multiple observations supporting a novel mechanism by which vFLIP promotes NF-κB signaling and latency. vFLIP is an established activator of NF-κB signaling and this activity is associated with viral latency. However, activation of NF-κB results in the expression of several negative regulators of the signaling pathway. Expression of one such negative regulator, A20, was shown to be induced by vFLIP. While NF-κB signaling is important for maintaining latency, prolonged NF-κB activation could contribute to an inflammatory phenotype. In fact, this is what occurs when negative regulators of NF-κB are either naturally or experimentally defective. Deficiencies in Itch ubiquitin ligase expression or function are associated with immune deficiencies and the Itch -/- knock out mouse displays an “itchy” phenotype for which this gene is named. A20 -/- mice also display a phenotype associated with inflammation and autoimmunity, exhibiting hypersensitivity to TNF and premature death. To establish and maintain a latent infection, vFLIP must activate NF-κB and signaling must be sustained without killing the host and to accomplish this, the virus must control negative regulators of NF-κB.

We observed interaction of vFLIP with the Itch and A20 ubiquitin editing complex. We previously reported that in the presence of RTA, Itch targets vFLIP for degradation. These observations suggest that vFLIP may be interacting with the Itch/A20 ubiquitin editing complex in latency and reactivation along with expression of RTA may be modulating the activity of this complex. We evaluated multiple known substrates of Itch via western blot and proteomic analysis and observed modest decreases in protein levels when RTA was expressed suggesting that RTA is altering Itch substrate stability. Our proteomics data revealed identification of 146 proteins with RTA dependent alterations in ubiquitination, however only two were known Itch or A20 substrates, suggesting that while RTA has a demonstrated effect on the cellular ubiquitome, the mechanism(s) governing this observation remains unclear. We reasoned that vFLIP interaction with the Itch A20 ubiquitin editing complex may function to promote NF-κB signaling, and expression of RTA abrogates signaling by inducing the degradation of vFLIP as well as other members of the complex.

vFLIP had no effect on Itch/A20 complex assembly, suggesting that vFLIP was not inhibiting protein complex formation as had been described with HTLV Tax[28]. We observed, through *in vitro* assay and through immunoprecipitation of RIPK1 conjugates, inhibition of A20 DUB activity by vFLIP. Detection of sustained K63 ubiquitinated RIP1 in the presence of vFLIP, suggests that A20 DUB activity is limited thereby allowing for sustained NF-ΚB signaling. We identified a SIM in vFLIP and observed interaction with SUMO-1 and 2/3. We previously reported that this motif is required for activation of NF-ΚB, degradation of vFLIP by RTA and interaction with Itch. Taken together these data suggest that vFLIP interacts with the Itch/A20 complex via a SUMO dependent mechanism. We observed evidence of Itch SUMOylation and inhibition of global SUMOylation resulted in a dose dependent increase in infectious virus production.

## Methods

### Cell Line Maintenance and Transfection

Human Embryonic Kidney 293T (HEK 293T) cells were cultured in DMEM medium supplemented with 10% Fetal Bovine Serum and were grown at 5% CO_2_ at 37°C. Cells were transfected at 60-70% confluency using 1µg/mL polyethyleneimine (PEI) linear, MW∼25,000 (Polysciences, Inc. Cat#23966) at a ratio of 1µg plasmid DNA: 3µl PEI. After 5 min of incubation the mixture was added to the cells. 24h post transfection the media was changed and if appropriate, 2.5µM of MG132 was added. Vero rKSHV.294 cells containing a recombinant reporter KSHV clone were cultured in DMEM, 10% FBS, and 10 µg/ml G418 antibiotic. MSR-tet OFF 293T cells were cultured in DMEM and 10% FBS. iSLK cells were obtained from Jae Jung via Young Bong Choi. Cells were maintained in DMEM supplemented with 10% Fetal Bovine Serum, penn/strep/neomycin, and 1ug/ml puromycin and grown at 5% CO_2_ at 37°C. BAC16 and BAC16 vFLIP V22E were introduced into iSLK cells using Lipofectamine 3000 followed by selection with hygromycin B (1200ug/ml), puromycin (1ug/ml), and G418 (250ug/ml).

### Reagents, Plasmids, and Antibodies

The proteasome inhibitor MG132 (Boston Biochem) was used in this study. FLAG-A20, was provided by Ed Harhaj, FLAG-Itch was provided by Annie Angers (REF), and myc-vFLIP by Gary Hayward. The following primary antibodies were used: anti-RTA (G. Hayward), anti-cmyc (Millipore), anti-Itch (BD Transduction Laboratories), anti-Flag (Sigma-Aldrich), anti-A20 (BD Transduction Laboratories), and anti-GFP (Thermo Scientific). The secondary antibodies used were anti-mouse-HRP, anti-rabbit-HRP, and anti-mouse AP (Jackson ImmunoResearch).

### Immunoblot Analysis

Proteins were run on 12% Tris-Glycine or Any kD mini-PROTEAN Precast Gel (Biorad) with Tris-glycine running buffer. The proteins were then transferred to a PVDF membrane using semi-dry transfer system at 20V for 20 minutes. The membranes were blocked in 5% non-fat dry milk in PBS for one hour. Primary antibodies were diluted in with 2.5% non-fat dry milk at 1uL antibody: 1000uL milk and applied to the membranes. The membranes were incubated on a shaker at 4°C overnight and were washed in PBS with 0.1% Tween the following day. Secondary antibodies were applied to the membranes in 2.5% non-fat dry milk at 1uL antibody: 1000uL milk. The membranes were incubated at room temperature on a shaker for one hour and afterward were washed with PBS and 0.1% Tween. Proteins were visualized with the addition of ECL substrate and the detection of the luminescence on x-ray film or scanned by a Li-COR C-DiGit Blot Scanner.

### Immunoprecipitation

Transfected cells with appropriate constructs were harvested 48h post-transfection with PBS and centrifuged at 1500 rpm for 10 min. The PBS was removed and 1mL of lysis buffer with 10µl of a protease inhibitor cocktail kit (Thermo Scientific) were added to each cell pellet. When appropriate 12.5µL of 5 mM NEM was added to each cell pellet. Cell lysates were centrifuged at 10,000 rpm for 5 minutes to remove cell debris. The resulting supernatant was precleared with protein A/G PLUS-agarose (Santa Cruz) for 30 min at 4°C. The lysates were transferred to a new 1.5mL tube and protein concentrations were measured and normalized with a Pierce BCA protein assay kit. Approximately 50 µg of protein was transferred to new 1.5mL tube to serve as control lysate. 1μg of the appropriate primary antibody was added to the remaining cell lysate and incubated on a rotator overnight at 4°C. 25µL of protein A/G-agarose were added the following day for 1hr and washed 4x with RIPA lysis buffer. 50µl of 2X Laemmli Buffer were added and samples were boiled at 100°C for 10 min. Samples were visualized through immunoblot analysis as described above

### GST-Pull-down Assay

A 5mL culture of BL21 cells containing the indicated constructs (GST tagged SUMO 1, SUMO 2/3, wildtype vFLIP or VE-vFLIP) was used to inoculate 50 mL of LB Broth. After 1.5-2 hours, when the OD = 0.4-0.7, expression was induced with 0.1mM IPTG. Cells were spun down and stored at -20°C overnight. Pellets were resuspended in 1 mL of PBS with 10µL of a protease inhibitor cocktail and sonicated. The samples were spun down at 9700 RPM for 1 minute and the supernatant transferred to a new tube. GST-SUMO samples were added to glutathione-sepharose and incubated at room temperature for an hour. Purified wildtype vFLIP and VE-VFLIP was harvested from transfected 293T cells for use for the *in vitro* assay. Transfected cells with appropriate constructs were harvested 48h post-transfection with PBS and centrifuged at 1500 rpm for 10 min. The PBS was removed and 1mL of lysis buffer with 10µl of a protease inhibitor cocktail kit (Thermo Scientific) were added to each cell pellet and the cells sonicated 4 times for 10s. The samples were spun down at 9700 RPM for 1 minute and the supernatant added to a new 1.5mL tube and would be incubated with the SUMO and glutathione-sepharose.

### A20 purification

A20-FLAG was transfected into 293T cells, and purified following 48 hours incubation. Nine 10 cm dishes of A20 transfected cells were harvested in ice cold PBS, centrifuged, and lysed in 1 mL 50 mM HEPES, pH 7.4, 100 mM KAc, 5 mM MgAc2, 100 μg/mL digitonin, 1 mM DTT, 1X EDTA-free Complete protease inhibitor cocktail (Thermo Scientific) for 20 min on ice. The lysate was centrifuged to remove cell debris. Supernatant was incubated with 100 μL of anti-Flag affinity resin (Sigma) for 1-1.5 hr at 4°C. The resin was washed three times in 1 mL lysis buffer, three times in 1 mL 50 mM HEPES, pH 7.4, 400 mM KAc, 5mM MgAc2, 100 μg/mL digitonin, 1 mM DTT buffer, and three times in 50 mM HEPES,pH 7.4, 100 mM KAc, 5 mM MgAc2 buffer. Elutions were carried out with one volume of 0.2 mg/ml 3X-Flag peptide in the final wash buffer at room temperature for 30 min. Two sequential elutions were combined to form the final fraction.

### In vitro deubiquitination assay

V5-His tagged vFLIP was expressed in *E. coli* (BL21) and purified using Ni-NTA resin (ThermoFisher). A20-Flag was purified as previously described. Purified tetra-K63 ubiquitin was purchased from Boston Biochem. The following reagents were added to 20μl reactions where indicated: A20 (2μM), vFLIP (1μM, 5μM, 10μM), tetra-K63 Ub (500nM). Reactions were incubated at 37ºC for 2hrs followed by the addition of 4x Laemmli loading buffer. Reactions were analyzed by SDS PAGE followed by immunoblot.

### Construction of BAC16 vFLIP V22E

Mutant BAC16 was constructed using the protocol of Tischer et al using BAC16 in GS1783 generously provided by Jae Jung[29–31]Briefly, using PCR (primers in Primers Table), a Kanamycin resistance gene containing an I-SceI restriction enzyme site flanked by regions from the KSHV genome containing the mutation of interest, was amplified. The kanamycin cassette was amplified from pEPkan-S and integrated into BAC16 using Red recombination. An overnight culture of GS1783, containing BAC16 and Red recombinase, was used to inoculate a fresh culture and grown to an OD of 0.5. The culture was transferred to a 42°C shaking water bath for 10 min. to induce Red recombinase expression followed by 10-20min incubation on ice. Bacteria were pelleted and washed 3x in ice cold water in preparation for electroporation. 100ng of kanamycin cassette were added to 50ul of cells in residual water and electroporated using the following parameters: 1.5kV, 25uF, 200 ohms, time constant=3.5-4.5 ms. 250ul SOC media was added to bacteria and incubated at 30°C for 1h for recovery. Cells were plated on LB agar containing kanamycin (20ug/ml) and incubated at 30°C for 24h. Kanamycin positive colonies were verified for insert by colony PCR using kanamycin specific primers. Following confirmation of kanamycin cassette, cells were cultured overnight at 30°C in LB containing kanamycin. Overnight cultures were used to inoculate LB containing chloramphenicol 15ug/ml (chlor resistance cassette is present in BAC16). To remove the kanamycin cassette, leaving the mutation behind, I-SceI expression was induced by adding arabinose to a final concentration of 1%. Cultured were incubated, shaking for 45 min at 30°C. To complete the removal of the kanamycin cassette, Red recombinase expression was again induced via incubation at 42°C for 10 minutes, followed by incubation at 30°C for 2h. 10x serial dilutions were plated on LB plates containing 15ug/ml chloramphenicol and 1% arabinose. Colonies were replicate plated on kanamycin to identify colonies that no longer contained the kanamycin resistance cassette. Chlor positive/kan negative clones were sequenced using vFLIP specific primers. Clones were further analyzed by sequencing (Illumina for full BAC16 and Sanger for repeats) and pulsed field gel electrophoresis.

### BAC16 sequencing

#### Library preparation, target enrichment, and Ion Torrent sequencing

Total DNA was quantified by Qubit 3.0 dsDNA HS Assay (Life Technologies). Libraries were manually prepared from 100 ng total DNA using the Ion Xpress Plus Fragment Library Kit (Life Technologies), protocol MAN0007044 Rev: C.0. Libraries were quantified and sized with Agilent Bioanalyzer 2100 High Sensitivity DNA Assay (Agilent Technologies) and pooled to 100pM final concentration. Templating and loading onto the Ion 530 Chip (Life Technologies) were automated on the Ion Chef (Life Technologies). Samples were sequenced on the Ion Torrent S5 (Life Technologies) with default parameters. Ion Torrent barcodes and adapter sequences are removed prior to output of FASTA sequence data files from the Ion Torrent S5 server.

#### Quality trimming, filtering, and read mapping

Reads were trimmed by low-quality base pairs (quality limit = 0.05), including 20 nucleotides at the 5′ terminal and 10 nucleotides at the 3’ terminal. All reads shorter than 50 nucleotides were filtered out. High -quality, trimmed reads were mapped to the KSHV genome (GenBank accession number NC_009333) using the map-to-reference tool on CLC Genomics Workbench v20.0.3 (Qiagen) with default parameters. A length fraction of 0.5 and a similarity fraction of 0.8 were selected and non-specific reads were ignored for better mapping accuracy.

#### PCR amplification of KSHV gap regions and Consensus sequence generation

Primer pairs tagged with the universal M13 primers were used to amplify the gap regions of the KSHV genome which are not typically covered by NGS technology[32]. PCR conditions used were previously described[32]. PCR products were Sanger Sequenced and mapped to the reference genome (NC_009333) along with the original NGS reads using the parameters described above. A consensus sequence was built on CLC Genomics Workbench (v20.0.3) based on quality score voting, with low coverage regions being defined as regions with a minimum of 0 read and a maximum of the total number of reads. Low coverage regions were filled from the reference sequence (NC_009333) and annotated as such.

### Quantitative real-time PCR

Cells were washed 3X with PBS and harvested by scraping and pelleting via centrifugation. RNA was extracted using the Promega SV Total RNA Extraction Kit. DNase-treated RNA was reverse transcribed using the RevertAid First Strand cDNA Synthesis Kit (Fermentas) according to manufacturer’s instructions. The resulting cDNA was used for qPCR performed on Biorad CFX Connect using Maxima SYBR Green/ROX qPCR Master Mix as per manufacturer’s specifications. RTA and RPS13 (housekeeping gene) were amplified using the primers listed in the primers table. Relative gene expression was calculated using the ΔC_T_ method using the RPS13 housekeeping gene for normalization. Error bars represent the standard deviation. Statistical analysis was conducted using GraphPad Prism 9 using an unpaired t test comparing WT-DOX to VE -DOX.

## Primers table

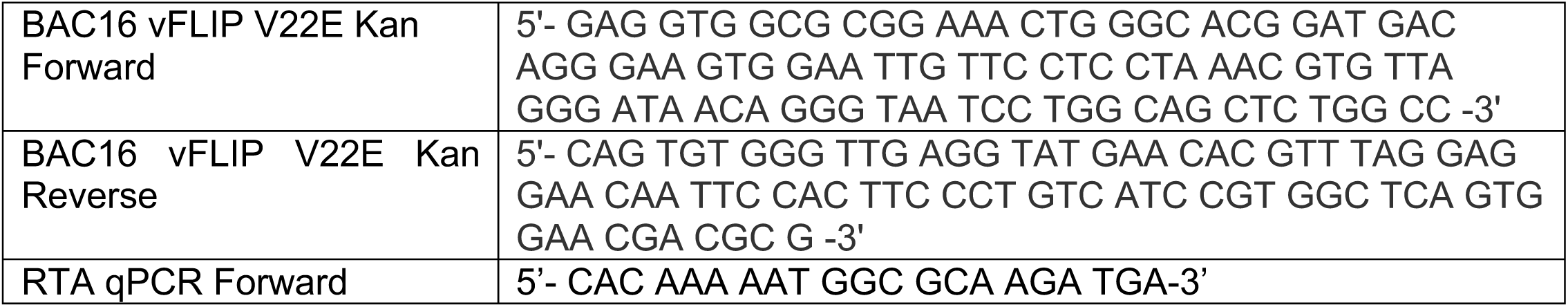
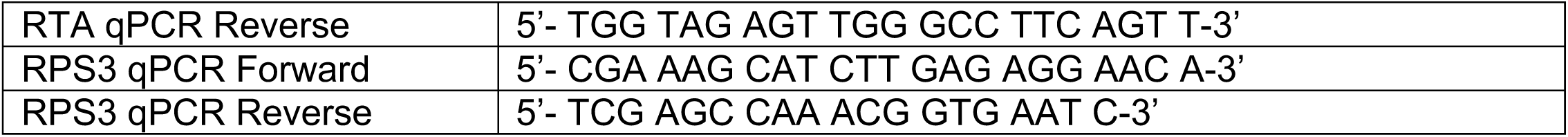

## Acknowledgements

This work was supported by funding to Elana S. Ehrlich from the Fisher Endowed Chair (Towson University) and NIH grant numbers GM118011, AI157907, and GM130382. Razia Moorad is supported by CA019014. Amerria Causey is supported by GM119970. We thank Ed Harhaj for the Itch and A20 expression vectors, Arie Admon and Aaron Ciechanover for hosting Elana S. Ehrlich’s sabbatical and providing support and advice for the ubiquitome proteomics, Gary Hayward for the RTA antibody, Jae Jung for the iSLK cells and the BAC16 in GS1783.

## References

1. Dittmer DP, Damania B. Kaposi sarcoma associated herpesvirus pathogenesis (KSHV) - An update. Current Opinion in Virology. 2013;3: 238–244. doi:10.1016/j.coviro.2013.05.012

2. Dissinger NJ, Damania B. Recent advances in understanding Kaposi’s sarcoma-associated herpesvirus. F1000Res. 2016;5: 740. doi:10.12688/f1000research.7612.1

3. Bower M, Nelson M, Young AM, Thirlwell C, Newsom-Davis T, Mandalia S, et al. Immune reconstitution inflammatory syndrome associated with Kaposi’s sarcoma. Journal of Clinical Oncology. 2005;23: 5224–5228. doi:10.1200/JCO.2005.14.597

4. Polizzotto MN, Uldrick TS, Hu D, Yarchoan R. Clinical manifestations of Kaposi sarcoma herpesvirus lytic activation: Multicentric Castleman disease (KSHV-MCD) and the KSHV inflammatory cytokine syndrome. Frontiers in Microbiology. 2012;3: 1–9. doi:10.3389/fmicb.2012.00073

5. Uldrick TS, Wang V, Mahony DO, Aleman K, Wyvill KM, Marshall V, et al. An Interleukin-6 Related Systemic Inflammatory Syndrome in Patients Co-Infected with Kaposi Sarcoma Associated Herpesvirus and HIV but without Multicentric Castleman Disease. 2010;1868: 350–358. doi:10.1086/654798 https://ntp.niehs.nih.gov/go/roc14 N (National TP. Report on Carcinogens, Fourteenth Edition. Research Triangle Park, NC: U.S.; 2016.

6. Cesarman E, Damania B, Krown SE, Martin J, Bower M, Whitby D. Kaposi sarcoma. Nature Reviews Disease Primers. 2019;5: 9. doi:10.1038/s41572-019-0060-9

7. Lee H-R, Li F, Choi UY, Yu HR, Aldrovandi GM, Feng P, et al. Deregulation of HDAC5 by Viral Interferon Regulatory Factor 3 Plays an Essential Role in Kaposis Sarcoma- Associated Herpesvirus-Induced Lymphangiogenesis. Palese Dirk Damania, Blossom PD, editor. mBio. 2018;9: e02217–17. doi:10.1128/mBio.02217-17

8. Thome M, Schneider P, Hofmann K, Fickenscher H, Meinl E, Neipel F, et al. Viral FLICE-inhibitory proteins (FLIPs) prevent apoptosis induced by death receptors. Nature. 1997;386: 517–521. doi:10.1038/386517a0

9. Chaudhary PM, Jasmin A, Eby MT, Hood L. Modulation of the NF-κB pathway by virally encoded Death Effector Domains-containing proteins. Oncogene. 1999;18: 5738–5746. doi:10.1038/sj.onc.1202976

10. Tolani B, Matta H, Gopalakrishnan R, Punj V, Chaudhary PM. NEMO Is Essential for Kaposi’s Sarcoma-Associated Herpesvirus-Encoded vFLIP K13-Induced Gene Expression and Protection against Death Receptor-Induced Cell Death, and Its N-Terminal 251 Residues Are Sufficient for This Process. Frueh K, editor. Journal of Virology. 2014;88: 6345 LP–6354. doi:10.1128/JVI.00028-14

11. Matta H, Mazzacurati L, Schamus S, Yang T, Sun Q, Chaudhary PM. Kaposi’s sarcoma-associated herpesvirus (KSHV) oncoprotein K13 bypasses TRAFs and directly interacts with the IkappaB kinase complex to selectively activate NF-kappaB without JNK activation. J Biol Chem. 2007;282: 24858–65. doi:10.1074/jbc.M700118200

12. Yang Z, Honda T, Ueda K. vFLIP upregulates IKKε leading to spindle morphology formation through RelA activation. Virology. 2018;522: 106–121. doi:10.1016/j.virol.2018.07.007

13. Hunte R, Alonso P, Thomas R, Bazile CA, Ramos JC, van der Weyden L, et al. CADM1 is essential for KSHV-encoded vGPCR-and vFLIP-mediated chronic NF-κB activation. PLOS Pathogens. 2018;14: e1006968.

14. Ding X, Xu J, Wang C, Feng Q, Wang Q, Yang Y, et al. Suppression of the SAP18/HDAC1 complex by targeting TRIM56 and Nanog is essential for oncogenic viral FLICE-inhibitory protein-induced acetylation of p65/RelA, NF-κB activation, and promotion of cell invasion and angiogenesis. Cell Death and Differentiation. 2019. doi:10.1038/s41418-018-0268-3

15. Brown HJ, Song MJ, Deng H, Wu T-T, Cheng G, Sun R. NF-κB Inhibits Gammaherpesvirus Lytic Replication. Journal of Virology. 2003;77: 8532 LP–8540. doi:10.1128/JVI.77.15.8532-8540.2003

16. Keller SA, Hernandez-Hopkins D, Vider J, Ponomarev V, Hyjek E, Schattner EJ, et al. NF-κB is essential for the progression of KSHV- and EBV-infected lymphomas in vivo. Blood. 2006;107: 3295 LP–3302. doi:10.1182/blood-2005-07-2730

17. Bravo Cruz AG, Damania B. In vivo models of oncoproteins encoded by Kaposi’s sarcoma-associated herpesvirus. Journal of Virology. 2019. doi:10.1128/jvi.01053-18

18. Nakayama R, Ueno Y, Ueda K, Honda T. Latent infection with Kaposi’s sarcoma-associated herpesvirus enhances retrotransposition of long interspersed element-1. Oncogene. 2019. doi:10.1038/s41388-019-0726-5

19. Sakakibara S, Espigol-Frigole G, Gasperini P, Uldrick TS, Yarchoan R, Tosato G. A20/TNFAIP3 inhibits NF-κB activation induced by the Kaposi’s sarcoma-associated herpesvirus vFLIP oncoprotein. Oncogene. 2012;32: 1223.

20. Ma A, Malynn B a. A20: linking a complex regulator of ubiquitylation to immunity and human disease. Nat Rev Immunol. 2012;12: 774–85. doi:10.1038/nri3313

21. Matta H, Gopalakrishnan R, Punj V, Yi H, Suo Y, Chaudhary PM. A20 is induced by Kaposi sarcoma-associated herpesvirus-encoded viral FLICE inhibitory protein (vFLIP) K13 and blocks K13-induced nuclear factor-kappaB in a negative feedback manner. J Biol Chem. 2011;286: 21555–64. doi:10.1074/jbc.M111.224048

22. Ehrlich ES, Chmura JC, Smith JC, Kalu NN, Hayward GS. KSHV RTA abolishes NFκB responsive gene expression during lytic reactivation by targeting vFLIP for degradation via the proteasome. PLoS One. 2014;9: e91359. doi:10.1371/journal.pone.0091359

23. Chmura JC, Herold K, Ruffin A, Atuobi T, Fabiyi Y, Mitchell AE, et al. The Itch ubiquitin ligase is required for KSHV RTA induced vFLIP degradation. Virology. 2017;501. doi:10.1016/j.virol.2016.11.016

24. Brown HJ, Song MJ, Deng H, Wu T. NF- κ B Inhibits Gammaherpesvirus Lytic Replication NF- B Inhibits Gammaherpesvirus Lytic Replication. J Virol. 2003;77: 8532–8540. doi:10.1128/JVI.77.15.8532

25. Grossmann C, Ganem D. Effects of NFkappaB activation on KSHV latency and lytic reactivation are complex and context-dependent. Virology. 2008;375: 94–102. doi:10.1016/j.virol.2007.12.044

26. DeCotiis JL, Ortiz NC, Vega BA, Lukac DM. An easily transfectable cell line that produces an infectious reporter virus for routine and robust quantitation of Kaposi’s sarcoma-associated herpesvirus reactivation. Journal of Virological Methods. 2017;247: 99–106. doi:10.1016/j.jviromet.2017.04.019

27. Shembade N, Pujari R, Harhaj NS, Abbott DW, Harhaj EW. The kinase IKKα inhibits activation of the transcription factor NF-κB by phosphorylating the regulatory molecule TAX1BP1. Nature Immunology. 2011;12: 834.

28. Brulois KF, Chang H, Lee AS-Y, Ensser A, Wong L-Y, Toth Z, et al. Construction and Manipulation of a New Kaposi’s Sarcoma-Associated Herpesvirus Bacterial Artificial Chromosome Clone. Journal of Virology. 2012;86: 9708–9720. doi:10.1128/jvi.01019-12

29. Tischer BK, von Einem J, Kaufer B, Osterrieder N. Two-step Red-mediated recombination for versatile high-efficiency markerless DNA manipulation in Escherichia coli. Biotechniques. 2006;40: 191–197. doi:10.2144/000112096

30. Jain V, Plaisance-Bonstaff K, Sangani R, Lanier C, Dolce A, Hu J, et al. A toolbox for Herpesvirus miRNA research: Construction of a complete set of KSHV miRNA Deletion Mutants. Viruses. 2016;8. doi:10.3390/V8020054

31. Moorad R, Juarez A, Landis JT, Pluta LJ, Perkins M, Cheves A, et al. Whole-genome sequencing of Kaposi sarcoma-associated herpesvirus (KSHV/HHV8) reveals evidence for two African lineages. Virology. 2022;568: 101–114. doi:10.1016/j.virol.2022.01.005

